# Metaplasticity in the ventral pallidum as a potential marker for the development of diet-induced obesity

**DOI:** 10.1101/2020.03.09.983114

**Authors:** Shani Gendelis, Dorrit Inbar, Kineret Inbar, Shanee Mesner, Yonatan M. Kupchik

**Affiliations:** Department of Medical Neurobiology, Faculty of Medicine, The Institute for Medical Research Israel-Canada (IMRIC), The Hebrew University of Jerusalem, Israel

## Abstract

A major driver of obesity is the increasing palatability of processed foods. Although reward circuits promote the consumption of palatable food their involvement in obesity remains unclear. The ventral pallidum (VP) is a key hub in the reward system that encodes the hedonic aspects of palatable food consumption and participates in various proposed feeding circuits. However, there is still no evidence for its involvement in developing diet-induced obesity. Here we examine, using male C57bl6/J mice and patch-clamp electrophysiology, how chronic high-fat-high-sugar (HFHS) diet changes the physiology of the VP and whether mice that gain the most weight differ in their VP physiology from others. We found that 10-12 weeks of HFHS diet hyperpolarized and decreased the firing rate of VP neurons without a major change in synaptic inhibitory input. Within the HFHS group, obesity-prone (OP, top 33% weight gainers) mice had a more hyperpolarized VP with longer latency to fire action potentials upon depolarization compared to obesity-resistant (OR, bottom 33% weight gainers) mice. OP mice also showed synaptic potentiation of inhibitory inputs both at the millisecond and minute ranges. Moreover, we found that the tendency to potentiate the inhibitory inputs to the VP might exist in overeating mice even before exposure to HFHS, thus making it a potential property of being an overeater. These data point to the VP as a critical player in obesity and suggest that hyperpolarized membrane potential of, and potentiated inhibitory inputs to, VP neurons may play a significant role in promoting overeating of palatable food.

**Significance statement:** In modern world, where highly-palatable food is readily available, overeating is often driven by motivational, rather than metabolic, needs. It is thus conceivable that reward circuits differ between obese and normal-weight individuals. But is such difference, if it exists, innate or develops with overeating? Here we reveal synaptic properties in the ventral pallidum, a central hub of reward circuits, that differ between mice that gain the most and the least weight when given unlimited access to highly-palatable food. We show that these synaptic differences exist also without exposure to palatable food, potentially making them innate properties that render some more susceptible than others to overeat. Thus, the propensity to overeat may have a strong innate component embedded in reward circuits.

## Introduction

Western calorie-rich diets are becoming more and more common in households. Consequently, obesity has become a world epidemic and a leading cause of preventable death worldwide (World Health Organization, 2003). Despite the drastic increase in obesity levels, the biological mechanisms driving this epidemic are still not understood and research is being conducted in many disciplines, including psychology, metabolism and neurobiology.

One main theory about the burst of the obesity epidemic in recent decades is that food has become during these years an increasingly strong reinforcer (Kenny, 2011). Various studies have indeed shown that not only do palatable foods act as reinforcers and can activate the reward system (de Macedo et al., 2016), but that direct activation of specific neurons in the reward system can drive by itself motivated feeding behavior (Nieh et al., 2015; Stuber and Wise, 2016; Friend et al., 2017). Such a strong link between the reward system and consumption of palatable foods poses the possibility that overeating that leads to obesity may involve pathologies in the reward system akin to those seen in models of addiction.

Research so far has indeed shown obesity-related changes both in the function of the reward system and in motivation-driven food seeking behavior. Thus, the expression level of the dopamine receptor D2/3 in the striatum is decreased in obese humans (Wang et al., 2001; Haltia et al., 2007; de Weijer et al., 2011) and rodents (Johnson and Kenny, 2010; Friend et al., 2017); opioid signaling in both the ventral tegmental area (VTA) (Vucetic et al., 2011; Martire et al., 2014), and the nucleus accumbens (NAc) (Alsiö et al., 2010; Vucetic et al., 2011) is altered in obese rodents; and the glutamatergic input to the NAc is potentiated in rats that develop diet-induced obesity (Brown et al., 2017; Oginsky and Ferrario, 2019), akin to the effect seen after withdrawal from cocaine (Gipson et al., 2013a, 2014) or nicotine (Gipson et al., 2013b, 2014). Possibly as a consequence of these changes in the reward system, rats and mice that gain the most weight in a diet-induced obesity model also show the highest motivation to obtain palatable food (Johnson and Kenny, 2010; Mancino et al., 2015; Vollbrecht et al., 2015; Brown et al., 2017; Derman and Ferrario, 2018; Inbar et al., 2019).

A key structure in the reward system, which is yet to be studied in the context of obesity, is the ventral pallidum (VP). The VP is a key junction between the reward system and a main hub for feeding behavior, the lateral hypothalamus (LH) (Stratford et al., 1999; Tripathi et al., 2013). It receives strong inhibitory input from the NAc (Zahm and Heimer, 1990; Kupchik et al., 2015) and in turn sends inhibitory projections to the LH and other targets (Root et al., 2015). Thus, it is not surprising that the VP has an important role in modulating feeding behavior (Cromwell and Berridge, 1993; Stratford et al., 1999; Shimura et al., 2006; Smith et al., 2009). Accordingly, application of GABA antagonists (Stratford et al., 1999; Covelo et al., 2014) or mu-opioid receptor agonists (Smith and Berridge, 2005) into the VP drive feeding, while fMRI studies in rodents, primates and humans show an increase in VP activity upon exposure to food or food cues (Hoch et al., 2013; Jiang et al., 2015; Royet et al., 2016; Kaskan et al., 2019). Nevertheless, whether the VP has a role in pathological overeating and obesity has not been studied. In this study we set to examine whether chronic consumption of HFHS diet alters the physiology of the VP and whether mice that are more prone to gain weight when HFHS food is freely available exhibit different VP physiology than those that are more resistant to gaining weight. Moreover, using behavioral and electrophysiological tools we test whether the differences in VP physiology between obesity-prone and resistant mice may be innate and thus possibly be part of the mechanism promoting overeating and weight gain.

## Materials and Methods

### Experimental subjects

Experimental subjects were naive C57bl6/J wildtype male mice weighing between 23g and 30g at the beginning of the experiment. A 12-hour reversed light/dark cycle was maintained, with the lights turned off at 8:00 am. Experimental procedures were conducted during dark hours. Mice were housed individually and nesting/enrichment material was available. Mice were given 7 days to acclimate before experimentation began. All experimental procedures were approved by the Authority for Biological and Biomedical Models in the Hebrew university.

### Diet-induced obesity model

After the acclimation period, all mice were put on a 4-week standard chow diet (Teklad Global 2018, 18% kcal fat; total density = 3.1 kcal g^−1^; Harlan Laboratories Inc., Indianapolis, Indiana), during which body weight and caloric intake were monitored and behavioral training was completed (see below). Then, mice that were put on the HFHS groups (n=24) were switched to a HFHS diet (D12451, 45% kcal fat; total density = 4.73 kcal g^−1^; Research Diets Inc.) for 10-12 weeks, while the control chow mice (n=15) continued to feed on the standard chow diet. At the end of these 10-12 weeks HFHS mice were switched back to standard chow for 2 additional weeks before being sacrificed for recordings. Body weight was determined twice per week (BJ-410C scales, Precisa, Dietikon, Switzerland) throughout the entire experiment. At the end of the diet-induced obesity protocol HFHS mice were divided according to their weight gain (relative to their weight on Day 1, when they were first exposed to the HFHS diet) into diet-induced obesity-prone (OP, top third weight gainers) and diet-induced obesity-resistant (OR, bottom third weight gainers) groups.

### Operant food self-administration protocol

Food self-administration was conducted in mouse operant boxes (MedAssociates, Fairfax VT, USA) as described before (Inbar et al., 2019). Briefly, mice were trained to press a lever in order to obtain a 20 mg precision pellet of chow food (F0071, 5.6% kcal from fat, total density = 3.6 kcal/g; Bioserv Inc., Frenchtown, NJ)) first on a fixed ratio 1 (FR1) schedule (each lever press resulted in a food pellet), and then on FR3 and FR5. Mice were eventually tested at the end of the HFHS diet on a progressive ratio task (where the number of lever presses required to obtain a pellet increased progressively). Progressive ratio schedule was 5, 9, 12, 15, 20, 25, 32, 40, 50, 62, 77, 95, 118, 145, 178, 219, 268, 328, 402, 492 and 603 lever presses per pellet. The test was terminated after 6 hours or if the mouse did not progress to the next level within one hour of achieving the previous one. During the test we monitored the number of lever presses on the active and inactive levers and the number of head entries into the food receptacle (detected by infrared beam). Operant chambers were located in sound-attenuating boxes to minimize external noises.

### Slice preparation

As described before (Inbar et al., 2020; Levi et al., 2020). Mice were anesthetized with 150 mg/kg Ketamine HCl and then decapitated. Sagittal slices (200 µm) of the VP were prepared (VT1200S Leica vibratome) and moved to vials containing artificial cerebrospinal fluid (aCSF (in mM): 126 NaCl, 1.4 NaH_2_PO_4_, 25 NaHCO_3_, 11 glucose, 1.2 MgCl_2_, 2.4 CaCl_2_, 2.5 KCl, 2.0 Na Pyruvate, 0.4 ascorbic acid, bubbled with 95% O_2_ and 5% CO_2_) and a mixture of 5 mM kynurenic acid and 100 mM MK-801. Slices were stored in room temperature (22-24 °C) until recording.

### In vitro whole cell recording

Recordings were performed at 32 °C (TC-344B, Warner Instrument Corporation). VP neurons were visualized using an Olympus BX51WI microscope and recorded from using glass pipettes (1.3-2 MΩ, World Precision Instruments) filled with internal solution (in mM: 68 KCl, 65 D-gluconic acid potassium salt, 7.5 HEPES potassium, 1 EGTA, 1.25 MgCl2, 10 NaCl, 2.0 MgATP, and 0.4 NaGTP, pH 7.2-7.3, 275mOsm). Excitatory synaptic transmission was blocked with CNQX (10 µM). Multiclamp 700B (Axon Instruments, Union City, CA) was used to record both membrane and action potentials and inhibitory postsynaptic currents (IPSCs) in whole cell configuration. Excitability and passive membrane properties were measured in current clamp mode while synaptic activity was measured in voltage clamp mode at a holding potential of −80 mV. Recordings were acquired at 10 kHz and filtered at 2 kHz using AxoGraph X software (AxoGraph Scientific, Sydney). To evoke IPSCs electrically, a bipolar stimulating electrode (FHC, Bowdoin, ME, USA) was placed ∼200-300 μm anterior of the cell to maximize chances of stimulating NAc afferents. The stimulation intensity chosen evoked 30% to 70% of maximal IPSC. Recordings were collected every 10 seconds. Series resistance (Rs) measured with a −2 mV hyperpolarizing step (10 ms) given with each stimulus and holding current were always monitored online. Recordings with unstable Rs or when Rs exceeded 20 MΩ were aborted.

### Current steps protocol

Neurons were recorded from in current clamp configuration (baseline membrane potential was adjusted to be approximately −50mV in all neurons). Five 500 ms-long depolarization current steps ranging from 0 pA to +80 pA (20 pA intervals) were applied, inter-step interval was 3 seconds. The 5-step protocol was repeated 5 times with 3 s between repetitions.

### High-frequency stimulation-induced plasticity measurements

Recorded in voltage clamp mode. Baseline evoked IPSCs were recorded at −80 mV for 3 min before applying the high-frequency stimulation (HFS) protocol. We used the same HFS protocol that was shown before to induce long-term plasticity (depression) in inhibitory synapses in the VP (Kupchik et al., 2014; Heinsbroek et al., 2017). The protocol consists of two trains separated by 20 s, each delivering 100 stimulations at 100 Hz. The amplitude of the pulses was identical to that inducing synaptic release when recording at −80 mV. During the HFS protocol, membrane potential was allowed to change freely (i.e. current clamp configuration). At the end of the second train of pulses recording was re-established at holding potential of −80 mV. Differences between groups were measured at the first minute after the HFS for post-tetanic potentiation and at minutes 13-19 for the long-term plasticity in all experiments.

### Paired-pulse ratio and coefficient of variation measurements

All experiments consisted of two consecutive stimulations with 50 ms interval. We measured the ratio between the amplitudes of the second and first pulse to calculate the paired-pulse ratio (PPR) and the coefficient of variation of the currents generated by the first of the two pulses. We repeated this at least 20 times for each condition.

### Data analysis

Statistics were performed using GraphPad Prism 8.2 (GraphPad Software Inc, San Diego, CA). Parametric statistics (student’s T-test, 1-way, 2-way or mixed effects ANOVA with Sidak’s multiple comparisons tests where appropriate) was used throughout except for Figs. 2A-B and 4A-B, where we used the Kolmogorov-Smirnov test. Lines in Figs. 3D, 4I and 5F-G were generated from simple non-parametric linear regression to the data. The ANCOVA test used in Figs. 2D-E, 4D-E, 5I compares the slope and intercept with Y-axis (called “elevation” here) of the linear regression of the datasets. If slopes are significantly different it is not possible to test whether the intercepts differ significantly. Thus, in Fig. 4E only the slope comparison is presented. If the slopes are not significantly different the intercepts are compared.

**Figure 1.**
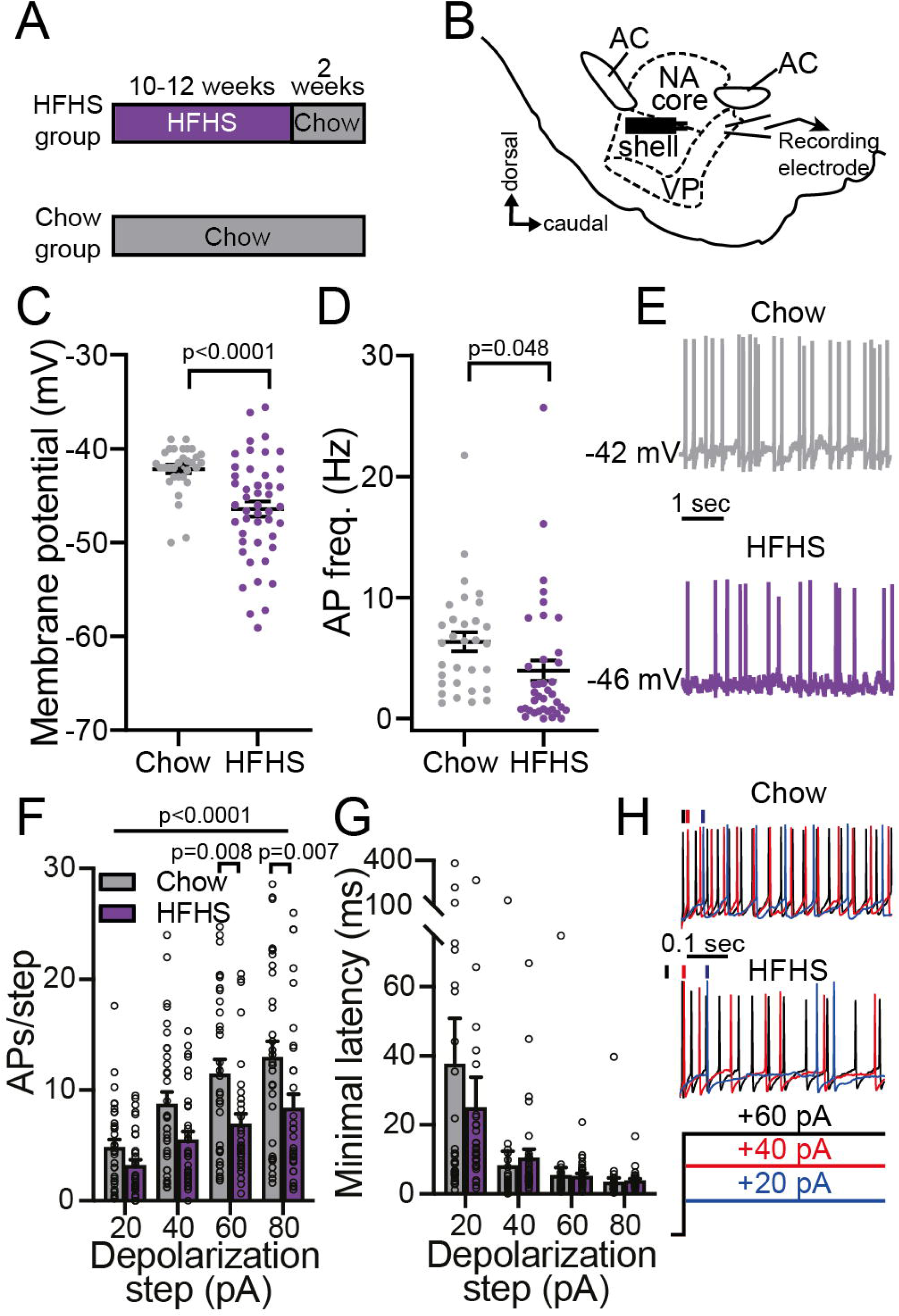
Chronic High-Fat-High-Sugar diet hyperpolarizes ventral pallidum neurons and inhibits their firing rate. (A) Experimental timeline. Mice in the HFHS group received a HFHS diet *ad libitum* in their home cages for 10-12 weeks and then returned to chow diet for 2 additional weeks before recordings. The control Chow group received regular chow during the same period of time. (B) Recording setup. VP neurons were recorded from in sagittal slices while activating incoming axons using a bipolar stimulating electrode placed ∼300 μm rostral to recorded cells. AC - anterior commissure; NA - nucleus accumbens. (C-E) VP neurons of HFHS mice had lower membrane potential (C) and lower action potential frequency (D) compared to chow mice (two-tailed unpaired t-tests). (E) Representative current clamp traces. (F-H) In a series of incrementing depolarizing steps the firing rate was significantly lower in VP neurons of HFHS mice compared with chow mice (F) (2-way ANOVA, main group effect F _(1,260)_ =23.04 p<0.0001, no group X step amplitude interaction; comparisons of individual bars using Sidak’s multiple comparisons test), but there was no difference in the minimal latency to the first action potential (G) (2-way ANOVA, p<0.5442). (H) Representative traces at 3 depolarizing steps. Data taken from 32-46 Cells in 14-21 mice per group.

**Figure 2.**
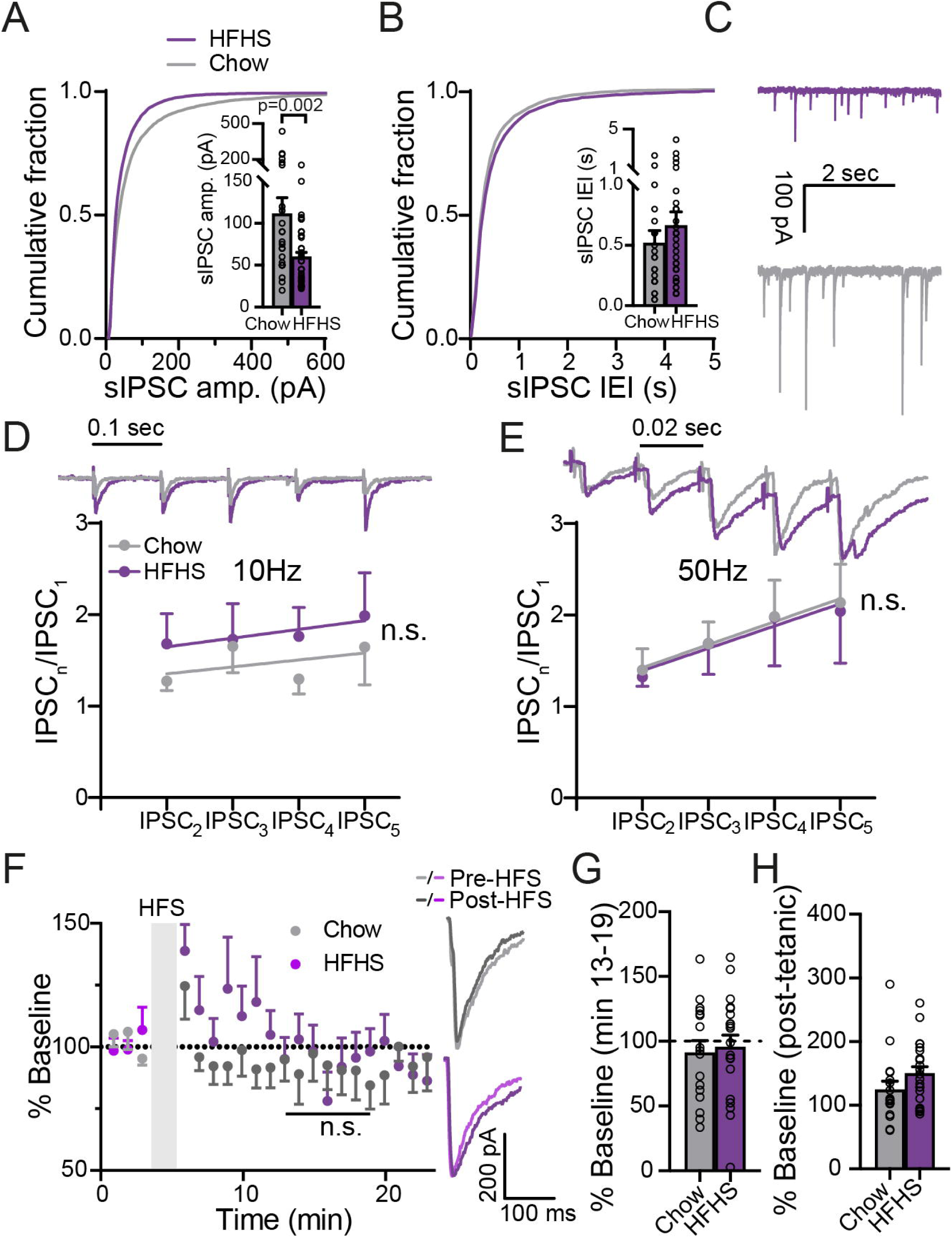
Chronic HFHS diet alters spontaneous but not evoked GABA neurotransmission in the ventral pallidum. (A) The cumulative probability plot of sIPSC amplitude is shifted to the left in HFHS mice, pointing to reduced sIPSC amplitude (Kolmogorov-Smirnov test, d=0.34, p=0.0002). Inset - median sIPSC amplitude was lower in HFHS mice (unpaired two-tailed t-test, t_63_=3.16). (B) HFHS diet did not affect the IEI of sIPSCs in the VP (Kolmogorov-Smirnov test, d=0.128, p=0.91). Inset-comparison of medians (unpaired two-tailed t-test, t_63_=0.84, p=0.41). (C) Representative sIPSC traces. (D-E) HFHS diet did not affect short-term plasticity induced by 5 consecutive stimulations at 10 Hz (D) or 50 Hz (E) (10 Hz 2-way ANCOVA, slope F=0.00875 p=0.9257, Elevation F=1.949 p=0.1641; 50 Hz 2-way ANCOVA, slope F=0.0009 p=0.9752, Elevation F=0.02642 p=0.8711). Insets representative traces synchronized with x axis. Stimulation artifacts were truncated. (F-G) The amplitude of evoked IPSCs after a high-frequency stimulation (HFS) did not differ between chow and HFHS mice (Mixed effects ANOVA, main group effect F _(1,35)_ = 0.016 p=0.901, interaction group X time F _(6,142)_ = 1.399 p=0.22; analysis performed on minutes 13-19), or from baseline (one sample two-tailed t-test on minutes 13-19, Chow - t_16_=0.96, p=0.35 HFHS - t_19_=0.47, p=0.65). Inset Representative traces. (H) Post-tetanic potentiation (PTP), measured at the first time point after the HFS, did not differ between groups (unpaired two-tailed t-test, t_37_=1.53, p=0.14). Data taken from 19-41 Cells in 13-24 mice per group.

**Figure 3.**
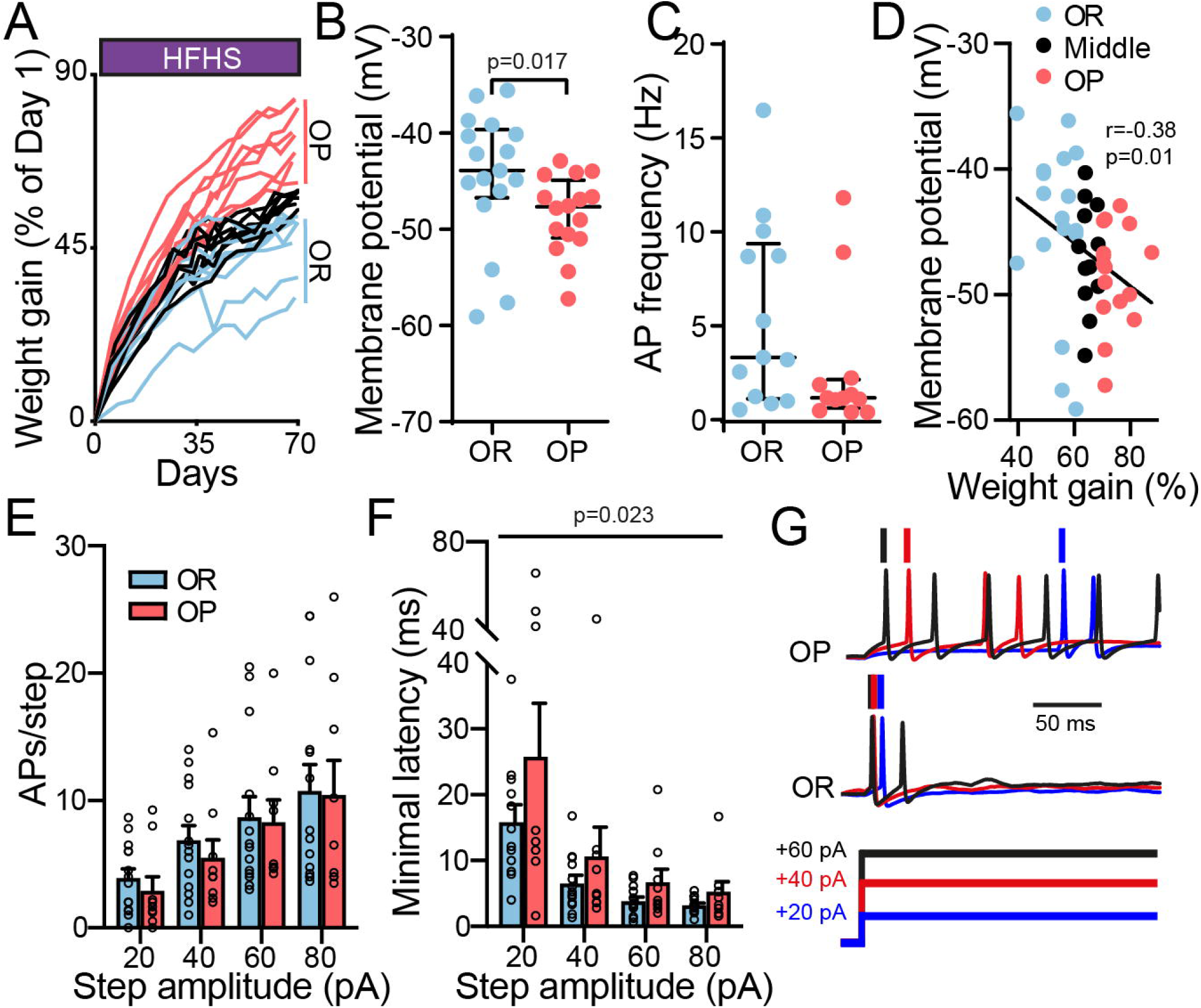
Ventral pallidum neurons of obesity-prone mice are more hyperpolarized and slower to fire compared with obesity-resistant mice. (A) Increase in body weight gain (expressed as % of 1st day) during 10-12 weeks of chronic HFHS diet for individual mice. Mice were split into obesity-resistant (OR, bottom 33% of weight gainers) and obesity-prone (OP, top 33% of weight gainers) groups according to final weight gain. (B) VP neurons of OP mice were more hyperpolarized than those of OR mice (unpaired two-tailed t-test, t_31_=1.99). (C) There was no significant difference in the baseline firing frequency of VP neurons between OP and OR mice (unpaired two-tailed t-test, t_31_=1.67, p=0.085). (D) VP membrane potential was negatively correlated with weight gain (Middle - middle 33% weight gainers). Correlation was assessed using non-parametric Spearman correlation. (E-F) The firing rates during incrementing depolarization steps did not differ between OP and OR mice (2-way ANOVA, main group effect F _(1,84)_ =0.4323, p=0.5126, interaction group X step amplitude F_(3,84)_=0.05, p=0.99), but the minimal latency to the 1st action potential was longer in HFHS mice (2-way ANOVA, main group effect F _(1,78)_ =5.37 p=0.0231, interaction group X step amplitude F_(3,84)_=0.73, p=0.54), possibly reflecting decreased excitability. (G) Representative traces. Data taken from 9-16 cells in 7 mice per group.

**Figure 4.**
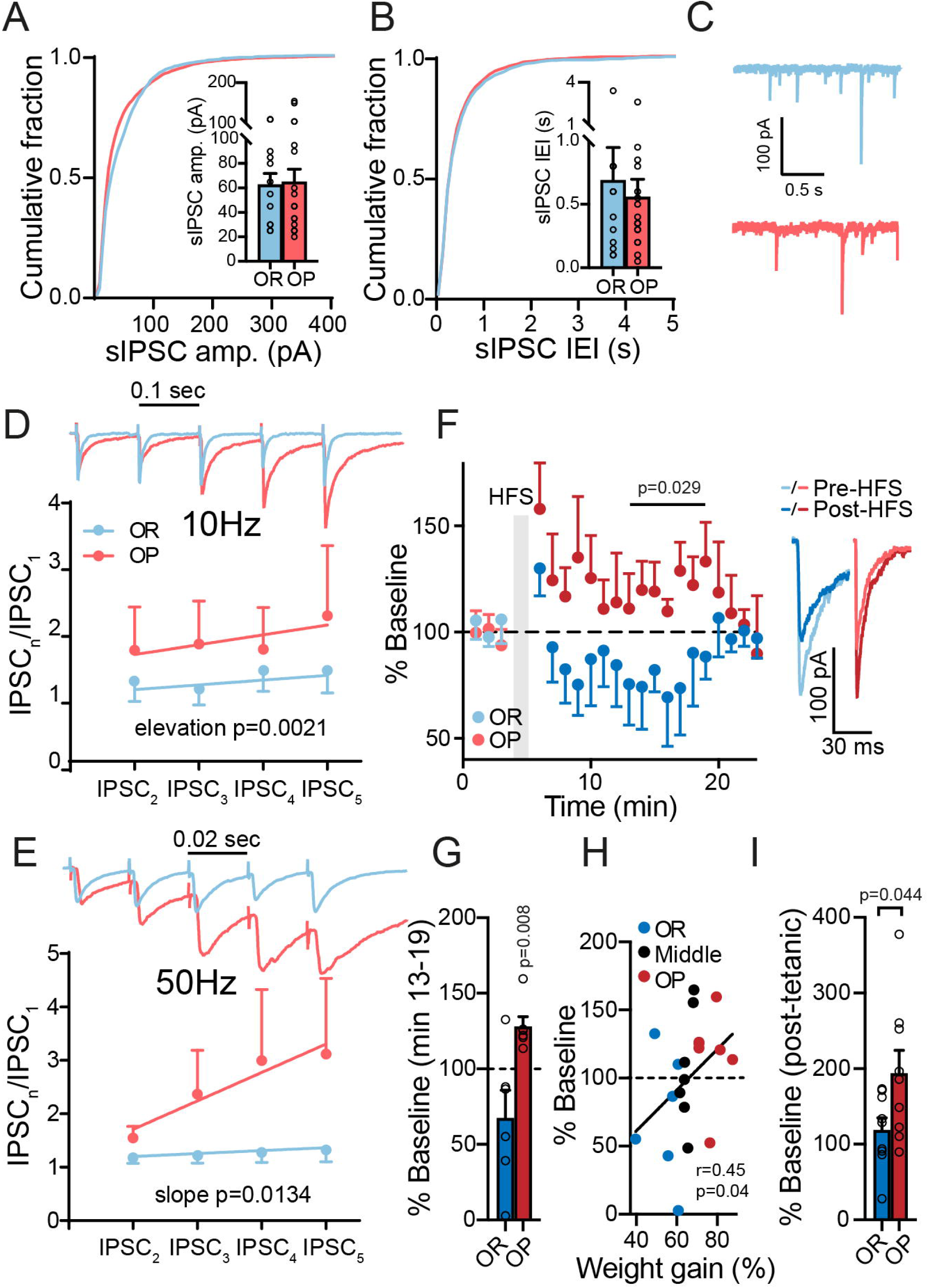
Obesity-prone and obesity-resistant mice show different synaptic plasticity in their GABA input to the VP. (A-C) sIPSC amplitude (A) and IEI (B) were similar between OP and OR mice (Kolmogorov-Smirnov tests - d=0.203, p=0.08 for amplitude and d=0.128, p=0.91 for IEI). Insets - comparison of medians (unpaired two-tailed t-tests, t_27_=0.15, p=0.88 for amplitude, t_27_=0.48, p=0.64 for IEI). (C) Representative sIPSC traces. (D-E) Five consecutive pulses at 10 Hz (D) or 50 Hz (E) generated short-term potentiation of IPSCs in OP but not OR mice (10 Hz 2-way ANCOVA, slope F_(1,4)_=0.5546 p=0.4978, elevation F_(1,4)_=34.32 p=0.0021; 50Hz 2-way ANCOVA, slope F_(1,4)_=17.78 p=0.0134). Insets representative traces synchronized with x axis. Stimulation artifacts were truncated. (F-G) HFS protocol given to inhibitory inputs to the VP generated a transient potentiation in OP mice but not in OR mice (Mixed effects ANOVA, main group effect comparing minutes 13-19, F _(1,12)_ = 5.29 p=0.029, interaction group X time F _(6,38)_ = 0.36 p=0.90; when comparing minutes 13-19 to 100% of baseline with one-sample two-tailed t-test - OP - t_5_=4.25, p=0.008; OR - t_5_=1.77, p=0.14). (F) Inset – representative traces. (H) HFS-induced plasticity in VP neurons was positively correlated with weight gain (Middle - middle 33% weight gainers), going from depression in OR mice to potentiation in OP mice. Correlation was assessed using non-parametric Spearman correlation. (I) Post-tetanic potentiation (PTP), measured at the first time point after the HFS, was stronger in OP mice (unpaired two-tailed t-test, t_16_=2.18, p=0.044). Data taken from 8-17 cells in 7 mice per group.

**Figure 5.**
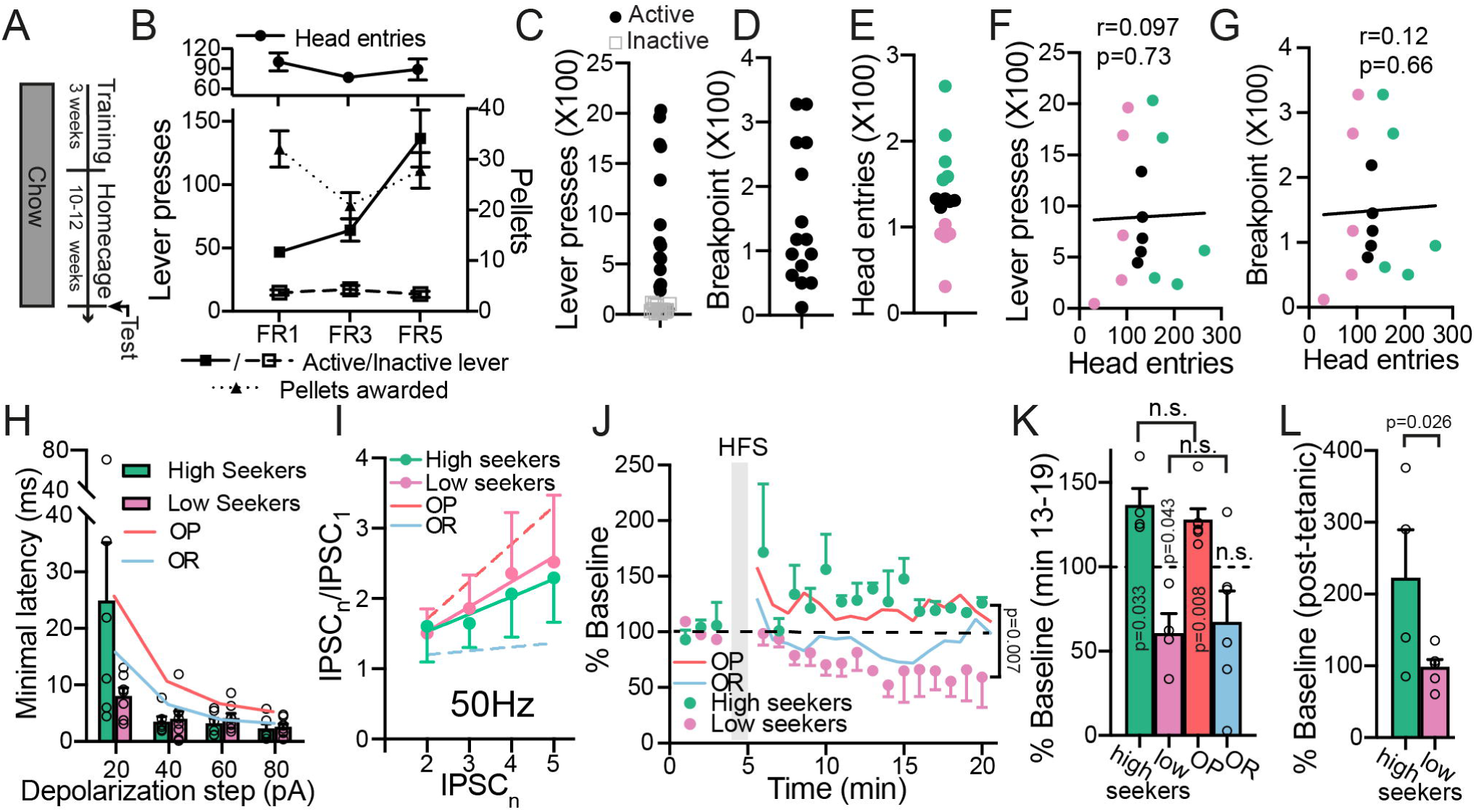
Differences in synaptic plasticity seen between OP and OR mice exist also between high and low food seekers without exposure to HFHS diet. (A) Experimental protocol. Mice were first trained to press a lever for 20 mg chow precision pellets. After 3 weeks of training mice were left in their home cages and fed on chow for 10-12 weeks. (B) Active and inactive lever pressing, pellets rewarded and head entries of mice during training. (C-E) After the HFHS diet mice were tested in the progressive ratio paradigm and lever pressing (C), breakpoint (D) and head entries (E) were recorded. Mice were split according to the head entries to the food receptacle (E) into “high seekers” (top third) and “low-seekers” (bottom third). (F-G) There was no correlation between head entries and lever presses (F) or breakpoint (G). Correlations were assessed using non-parametric Spearman correlation. (H) There was no difference in the minimal delay to the first action potential in a series of depolarizing steps (from +20 pA to + 80 pA) between high and low seekers (Mixed effects analysis, no main group effect, F_(1,12)_=1.9, p=0.19, interaction group X step amplitude F_(3,35)_=3.73, p=0.02). (I) There was no difference in short-term plasticity between high and low seekers (5 stimulations at 50 Hz; Two-way ANCOVA, slope F_(1,4)_=0.075 p=0.785, Elevation F_(1,4)_=0.140 p=0.71). (J-K) HFS protocol potentiated eIPSCs in the high-seekers and depressed eIPSCs in the low-seekers (Mixed effects ANOVA comparing minutes 13-19, main group effect F (1,7) = 16.13 p= 0.007, interaction group X time F _(7,22)_ = 0.72 p=0.65; comparing minutes 13-19 to 100% of baseline - one-sample t-test, t_3_=3.75 p=0.033 and t_3_=3.38 p=0.043 for high and low seekers, respectively). These differences in HFS-induced plasticity between high and low seekers were similar in amplitude to the differences seen between OP and OR mice (depicted as lines) after HFHS diet (Mixed effects ANOVA, no group effect when comparing OP mice to high-seekers (F_(1,9)_=0.7375 p=0.4128) or OR mice to low-seekers (F_(1,12)_=0.2007 p=0.6622); p>0.05 using two-tailed unpaired t-test when comparing minutes 13-19 in high seekers to OP mice or low seekers to OR mice). (L) Post-tetanic potentiation (PTP), measured at the first time point after the HFS, was stronger in high seekers compared to low seekers (unpaired one-tailed t-test, t_8_=2.28, p=0.026). Data taken from 4-9 cells in 5 mice per group.

## Results

### High Fat High Sugar diet hyperpolarizes membrane potential and reduces firing rate in VP neurons

In order to establish the long-lasting influence of chronic consumption of highly palatable food on the VP of mice, we exposed two groups of mice to different diet protocols. The first group received high-fat-high-sugar (HFHS) diet *ad libitum* in the home cage for 10-12 weeks followed by two weeks of standard chow diet. The second group received standard chow for a similar time period (Fig. 1A).

We then examined, using whole-cell patch clamp electrophysiology, whether the long exposure to HFHS had affected the cellular properties of VP neurons (Fig. 1B). We found that both resting membrane potential (Fig. 1C) and action potential firing frequency (Fig. 1D-E) were significantly lower in the VP of HFHS mice compared to the Chow mice. Moreover, we found that when depolarized to different levels, VP neurons of HFHS mice fired significantly less than VP neurons of Chow mice, an effect that was most robust at the strongest depolarization steps (+60 and +80 pA) (Fig. 1F,H). The minimal latency to the first action potential in a depolarizing step (Fig. 1G) did not differ between groups. Examination of the shape of the action potentials revealed a wider waveform in the HFHS group but other parameters did not differ between groups (Table 1). These data demonstrate that chronic HFHS consumption changes the physiology of VP neurons, rendering them more hyperpolarized and less active, even two weeks after the last exposure to HFHS.

**Table 1.** Action potential and passive membrane properties

|  | HFHS mice | Chow mice | P value | OP mice | OR mice | P value |
| --- | --- | --- | --- | --- | --- | --- |
| AP Amplitude | 52.06±12.21<br>(n=34) | 46.14±13.79<br>(n=25) | P=0.0868 | 48.52±7.46<br>(n=10) | 55.82±14.92<br>(n=15) | P=0.1526 |
| AP Half width | 1.706±0.829<br>(n=34) | 1.258±0.581<br>(n=26) | P=0.0226 | 1.739±0.676<br>(n=10) | 1.359±0.484<br>(n=15) | P=0.1140 |
| AP rise time | 1.784±0.915<br>(n=34) | 1.181±0.591<br>(n=26) | P=0.005 | 2.103±1.083<br>(n=10) | 1.308±0.516<br>(n=15) | *P=0.0213 |
| AP decay | 1.714±0.861<br>(n=34) | 1.324±0.747<br>(n=26) | P=0.071 | 1.620±0.7162<br>(n=10) | 1.443±0.583<br>(n=15) | P=0.5036 |
| AHP peak | -12.9±4.112<br>(n=29) | -14.2±3.537<br>(n=23) | P=0.2367 | -13.65±2.906<br>(n=8) | -12.25±5.08<br>(n=14) | P=0.4867 |
| AHP Half width | 8.281±4.393<br>(n=17) | 7.46±2.648<br>(n=9) | P=0.6141 | 10.29±3.307<br>(n=5) | 7.878±5.006<br>(n=8) | P=0.3631 |
| Series Resistance | 6.457±2.302<br>(39) | 6.735±3.257<br>(n=26) | P=0.7114 | 6.099±1.968<br>(n=9) | 6.583±2.999<br>(n=12) | P=0.521 |
| Capacitance | 7.475±2.009<br>(n=26) | 7.816±2.392<br>(n=12) | P=0.6502 | 8.077±1.654<br>(n=10) | 7.794±1.758<br>(n=8) | P=0.7301 |
| Membrane resistance | 366.8±170.9<br>(n=26) | 261.6±58.39<br>(n=12) | *P=0.0463 | 282.8±154<br>(n=10) | 347±138.5<br>(n=8) | P=0.3722 |
| AP – Action potential; AHP – After Hyperpolarization |  |  |  |  |  |  |

### High fat high sugar diet effect on GABA neurotransmission in the VP

The changes in cellular physiology induced by the chronic HFHS diet may be accompanied by changes in synaptic neurotransmission in the VP. To test for that we compared evoked and spontaneous inhibitory postsynaptic currents (eIPSCs and sIPSCs, respectively) between the HFHS and the Chow groups. When examining the amplitude and frequency of sIPSCs, we found that chronic HFHS diet significantly decreased the amplitude (Fig. 2A,C) but not the frequency (Fig. 2B,C) of sIPSCs. This reflects changes in GABA neurotransmission in the VP. Changes in the amplitude of spontaneous events are sometimes interpreted as reflecting postsynaptic mechanisms, but as these include action-potential-induced events, we think that this claim cannot be made for our data. Nevertheless, the changes we observed are less likely to be of presynaptic origin as we did not find an effect on the coefficient of variation (CV) and paired-pulse ratio (PPR) of eIPSCs, two indicators of changes in the probability of synaptic release (Faber and Korn, 1991; Berninger et al., 1999; Schinder et al., 2000) (Table 2).

**Table 2.** Coefficient of variation (CV) and paired-pulse ratio (PPR) of evoked IPSCs at baseline and after the HFS

|  | HFHS mice | Chow mice | P value | OP mice | OR mice | P value |
| --- | --- | --- | --- | --- | --- | --- |
| CV | 39.77±10.33<br>(n=24) | 37.56±10.6<br>(n=15) | P=0.5225 | 38.71±9.892<br>(n=5) | 33.51±7.574<br>(n=8) | P=0.3062 |
| CV after HFS | 44.45±9.486<br>(n=23) | 44.26±10.45<br>(n=15) | P=0.9541 | 44.06±7.611<br>(n=5) | 43.08±44.24<br>(n=8) | P=0.8672 |
| PPR | 1.302±0.568<br>(n=24) | 1.445±1.233<br>(n=19) | P=0.6163 | 1.364±0.812<br>(n=8) | 1.376±0.559<br>(n=8) | P=0.9731 |
| PPR after HFS | 1.818±1.228<br>(n=16) | 1.797±1.14<br>(n=16) | P=0.9618 | 1.245±0.535<br>(n=4) | 2.033±1.293<br>(n=5) | P=0.2954 |

To examine whether chronic HFHS diet changes also evoked GABA neurotransmission in the VP we examined the generation of synaptic plasticity in the millisecond (short-term plasticity) and the minute (long-term plasticity) range. Short-term plasticity was induced by applying 5 identical pulses at 1 Hz, 10 Hz, or 50 Hz and the change in eIPSC during the protocol was calculated by dividing each eIPSC by the first one. Our data show that the HFHS diet did not affect short-term plasticity in the VP in either of the frequencies (Fig. 2D,E; data for 1 Hz not shown). To induce long-term plasticity, we applied a high frequency stimulation (HFS) protocol that we and others used before (Kupchik et al., 2014; Creed et al., 2016) to elicit transient LTD in the VP. In contrast to previous findings, this protocol did not elicit plasticity or affect the CV or PPR parameters in the chow or HFHS groups (Fig. 2F-G, Table 2) (the lack of plasticity at the group level does not reflect lack of plasticity at the single cell level. As will be shown below, some cells showed depression while others showed potentiation). The HFS protocol did generate post-tetanic potentiation in both groups (measured as the difference from baseline of the eIPSCs during the first minute after the HFS) but without difference between them (Fig. 2H). Overall, the data suggest that chronic HFHS diet impairs spontaneous neurotransmission possibly through a postsynaptic mechanism but does not affect the short- and long-term plasticity of evoked GABA neurotransmission.

### Obesity-prone mice show hyperpolarized membrane potential and delayed firing in the VP compared to obesity-resistant mice

Our data so far show that chronic exposure to HFHS food changes the physiology of the VP. Our next question was whether within the group of mice exposed to HFHS diet, those that gained the most weight (top 33% weight-gainers relative to day 1 of the HFHS diet, called obesity-prone, OP) showed different physiology in the VP compared with those who gained the least weight (bottom 33% weight gainers, called obesity-resistant, OR) (Fig. 3A). Previous studies showed that not only does weight gain correlate with addictive-like behaviors towards palatable food (Vollbrecht et al., 2015; Brown et al., 2017; Inbar et al., 2019; Oginsky and Ferrario, 2019), it also correlates with synaptic changes in the NAc (Brown et al., 2017; Oginsky and Ferrario, 2019) - OP rats show stronger excitatory input to the NAc compared to OR rats. As the VP is a major target of the NAc, we expect that the physiology of the VP will also differ between OP and OR mice.

Membrane potential of VP neurons was significantly more hyperpolarized in OP mice compared with OR mice (Fig. 3B), although action potential frequency did not differ between OP and OR mice (Fig. 3C). Membrane potential (Fig. 3D), but not firing frequency (Table 3), also showed an inverse correlation with weight gain across the entire HFHS population. Applying depolarizing steps of increasing amplitudes from baseline membrane potential induced a comparable number of action potentials in both groups (Fig. 3E), but the minimal latency to the first action potential, a measure inversely correlated with cellular excitability (Alexander et al., 2019), was significantly longer in the OP mice (Fig. 3F-G). Both these parameters did not correlate with weight gain when examining the entire group of mice (Table 3). Overall, these data show that VP neurons of OP mice are more hyperpolarized and are slower to fire compared with those of OR mice.

**Table 3.** Correlation of weight gain with various neurophysiological parameters in the VP

|  | AP frequency | APs/+80 pA step | Minimal latency to 1<sup>st</sup> AP (+20 pA step) | sIPSC amplitude | sIPSC frequency (IEI) | IPSC<sub>5</sub>\IPSC<sub>1</sub> (50HZ) |
| --- | --- | --- | --- | --- | --- | --- |
| Weight gain | p=0.06<br>r=-0.31 | p=0.56<br>r=-0.1 | p=0.62<br>r=-0.1 | p=0.4<br>r=0.14 | p=0.87<br>r=0.03 | p=0.17<br>r=0.25 |
| r – Spearman's r, non-parametric Spearman correlation |  |  |  |  |  |  |

### Obesity-prone mice show altered synaptic plasticity in GABA input to the VP

To evaluate whether GABA synaptic transmission in the VP is also different between OP and OR mice we compared spontaneous and evoked IPSCs with the same tools as in Fig. 2. An examination of the amplitude and frequency of the inhibitory spontaneous events in the VP showed that these parameters were comparable between OP and OR mice (Fig. 4A-C) and did not correlate with weight gain in the entire HFHS population (Table 3). There was also no difference in the CV or PPR of eIPSCs between the groups (Table 1). The ability of VP neurons to show short-term plasticity, on the other hand, was different between OP and OR mice. Application of 5 consecutive stimulations at 10 or 50 Hz (but not at 1 Hz, data not shown) induced a strong potentiation of the IPSC in OP mice but not in OR mice (Fig. 4D-E), even though the amplitudes of the first IPSCs were similar (data not shown). This could mean that in OP mice, but not OR mice, fast repeated activation of inhibitory inputs to the VP may result in stronger inhibition that builds up with activity. The short-term plasticity showed a positive correlation with weight gain but this correlation did not reach significance (Table 3).

VP neurons of OP and OR mice also showed a striking difference in their longer-term response to a high-frequency stimulation (HFS) (Fig. 4F). Application of the HFS protocol (Kupchik et al., 2014) in VP neurons of OR mice generated a non-significant transient decrease in the average IPSC amplitude to 67.4±45% of baseline (Fig. 4F-G). In contrast, the HFS protocol induced a transient potentiation in VP cells of OP mice that lasted for ∼15 minutes (Fig. 4F-G). In fact, the direction and amplitude of the HFS-induced plasticity correlated with weight gain after chronic HFHS across all mice, transforming from depression in OR mice to potentiation in OP mice (Fig. 4H). In addition, the HFS generated a more robust post-tetanic potentiation in OP mice (Fig. 4I), again pointing to a tendency to strengthen inhibitory input in these mice after tetanic stimulation. Note the similarity in the results for the long and the short-term plasticity protocols-in both cases the inhibitory input to the VP of OP mice was potentiated while the input to the VP of OR mice did not change or even showed depression in some cells. Overall, the data in Fig. 4 show a difference in synaptic plasticity of inhibitory input to the VP between OP and OR mice, with a tendency towards potentiation of these inputs in OP mice.

### Differences in HFS-induced plasticity between OP and OR mice may be innate

In this and previous works, we and others showed various synaptic and physiological parameters in the NAc and the VP that differ between OP and OR animals (Brown et al., 2017; Oginsky and Ferrario, 2019). For obvious reasons, slice recordings in all these studies could only be performed *after* the HFHS diet and splitting mice to OP and OR groups. This technical hurdle left one of the most important questions unanswered - are these synaptic differences between OP and OR mice the cause or the consequence of obesity?

In a recent study (Inbar et al., 2019) we showed that in a progressive ratio task, where the reward is a small pellet of HFHS food, head entries to the food receptacle are higher in OP mice compared to OR mice even before the beginning of the diet. Based on this, we hypothesize here that if a physiological difference in the VP between OP and OR mice is innate, it may also exist without exposure to HFHS diet between mice that show the highest and the lowest rates of head entries in a progressive ratio task.

When choosing the physiological parameters to test, we specifically looked for parameters that on the one hand were not affected by the mere exposure to HFHS diet but on the other hand did differ between OP and OR mice after prolonged HFHS consumption. This is because we looked for innate differences between OP and OR mice. Thus, any difference driven by the HFHS diet itself (Figs. 1-2) cannot be confirmed as being innate. Based on these a-priori conditions, we examined the only physiological measures that fulfilled the requirements – the latency to fire upon depolarization and synaptic plasticity.

To examine whether the difference between OP and OR mice in either of the physiological measures listed above is innate, we trained a chow group in a progressive ratio task. Briefly, mice were first trained to press a lever in order to receive a 20 mg pellet of chow food (see Materials and Methods for details) and then were left in their home cages for 10-12 weeks with regular chow diet (Fig. 5A). Then, their performance (head entries into the food receptacle and lever presses) was measured during a progressive ratio task - the number of lever presses required for the delivery of a 20 mg pellet increased progressively until mice did not reach the next requirement (Figs. 5B-D). We then split the mice to those who showed the highest (top 33%, “high seekers”) and lowest (bottom 33%, “low seekers”) number of head entries (Fig. 5E). Note that there was no correlation between the head entries and the lever presses or breakpoint (Fig. 5F-G), confirming our previous finding that head entries, but not operant food seeking, may reflect better an innate property of OP mice (Inbar et al., 2019).

The latency to fire upon depolarization and the short-term plasticity of evoked GABA release showed no difference between high- and low-seekers (Fig. 5H-I), unlike the results in OP and OR mice (Figs. 3-4). This suggests that these differences may be a consequence of the HFHS diet rather than innate differences. In contrast, application of the HFS protocol to induce long-term plasticity showed a significant difference between high-seekers and low-seekers - it depressed IPSCs in the VP of low-seekers but potentiated IPSCs in high-seekers (Fig. 5J-K). This is strikingly similar to the difference in HFS-induced plasticity between OP and OR mice (Fig. 4F-H, Fig. 5K), although note that the plasticity seen in high and low seekers seem to last longer and not be as transient as in OP and OR mice. Moreover, the HFS-induced post-tetanic potentiation in high-seekers was stronger than in low-seekers (Fig. 5L), similar to the difference between OP and OR mice (Fig. 4I). Thus, the nature of HFS-induced plasticity in inhibitory inputs to the VP may be an innate property that could determine the level of weight gain in a chronic highly-palatable diet, and possibly be part of the cause for the extreme weight gain in OP mice.

## Discussion

In this work we showed a tight connection between the physiology of the VP and diet-induced obesity. First, we showed that chronic HFHS diet changed both cellular physiological properties and inhibitory synaptic input in VP neurons. Accordingly, VP neurons in HFHS-fed mice were more hyperpolarized and fired less action potentials (Fig. 1), and showed altered spontaneous GABAergic input (Fig. 2). We also showed that the physiology of VP neurons differed between OP and OR mice. VP neurons of OP mice were more hyperpolarized and slower to fire upon depolarization compared to those of OR mice (Fig. 3) and their inhibitory input tended to potentiate after protocols that induce plasticity in the millisecond and minute range (Fig. 4). Lastly, we were able to identify the tendency to show potentiation of GABA inputs in the VP as a possible innate marker for excessive palatable food seeking and body weight gain. In a previous study we showed that OP mice seek for palatable food more intensely than OR mice even before being exposed to chronic HFHS diet (Inbar et al., 2019). Here we showed that “high seekers” of palatable food never before exposed to HFHS diet showed potentiation of GABA input akin to OP mice while “low seekers” showed depression comparable in amplitude to that seen in OR mice (Fig. 5). Overall, our data pose the VP as a central player in diet-induced obesity that may contribute to the differences in diet-induced weight gain between individuals.

### The ventral pallidum, feeding and obesity

To date there are no studies linking diet-induced obesity to physiological changes in the VP and only few show changes in the striatum in obese individuals. For example, the expression of the striatal inhibitory D2 dopamine receptor is decreased both in obese humans (Wang et al., 2001) and animal models of obesity (Johnson and Kenny, 2010; Friend et al., 2017). Also, the excitatory input to the NAc is stronger in obese rats (Brown et al., 2017; Oginsky and Ferrario, 2019). Both these findings point to a possible increase in the activity of NAc GABAergic inputs to the VP in obesity. This may imply stronger inhibitory input on the VP in obesity. In line with this hypothesis, our data show a tendency to potentiate GABAergic input to the VP of OP mice (Fig. 4).

Our data also links diet-induced obesity to a more hyperpolarized and slower-to-fire VP (Figs. 1,3). The activity of the VP is crucial in determining food consumption. Feeding is associated with increased activation in the VP (Shimura et al., 2006; Zhu et al., 2017) while reducing VP activity by various methods (Stratford et al., 1999; Shimura et al., 2006; Smith et al., 2009; Chang et al., 2017) inhibits food consumption. The VP also encodes the hedonic value of food (Tindell et al., 2006; Smith et al., 2009) - blocking GABA neurotransmission increases food and sucrose seeking, but not water or quinine consumption (Stratford et al., 1999; Smith and Berridge, 2005; Shimura et al., 2006) and devaluation of palatable food leads to decreased VP activity (Brownell and Walsh, 2018). It seems, thus, that our data contradicts the main current thinking correlating VP activity positively with food consumption. However, it is important to keep in mind that here we tested the *baseline* activity of the VP in a pathological condition and not its acute response during healthy feeding behavior.

Little is known about the role of the VP in obesity. Human studies show an inverse correlation between VP activity to unappetitive images and Body Mass Index (BMI) (Burger and Stice, 2014) and higher levels of the serotonin receptor 5HT4R in the VP and NAc in obese individuals (Haahr et al., 2012). Other studies show a decrease in pallidal grey matter and network efficiency in obese patients (Baek et al., 2017; Hamer and Batty, 2019), although in these studies the globus pallidus is included in the analyses. At this point it is hard to integrate our data, taken at the synaptic level, with present human studies to form a solid hypothesis on the role of the VP in obesity and further research is needed.

### Synaptic plasticity and the susceptibility to develop obesity or addiction

One of our main findings is that VP neurons of OP and OR mice show different plasticity patterns upon application of a HFS (Fig. 4). A similar difference is seen also between high and low food seekers never exposed to HFHS (Fig. 5). Thus, the pattern of HFS-induced synaptic plasticity may be an innate property that may drive some but not others to seek for palatable food and become obese. This finding can be paralleled to the differences in metaplasticity (the plasticity of synaptic plasticity (Abraham and Bear, 1996)) sometimes seen in animal models of addiction. The glutamatergic input to the NAc is a plastic input that can show both LTP and LTD in drug naive animals (Kombian and Malenka, 1994). Chronic exposure to cocaine, with or without subsequent withdrawal, impairs this plasticity - both LTD (Thomas et al., 2001; Martin et al., 2006; Moussawi et al., 2009; Kasanetz et al., 2010) and LTP (Moussawi et al., 2009) induced by experimental protocols *ex vivo* and *in vivo* in the NAc are abolished. Similarly we have shown loss of plasticity in the VP after withdrawal from cocaine (Kupchik et al., 2014). The impaired plasticity in the NAc seems to not only be a general phenomenon in addiction but to inversely correlate with the susceptibility to develop addiction (Kasanetz et al., 2010). Thus, rats that show the highest level of cocaine self-administration do not show LTD while rats with minimal cocaine seeking behavior have intact LTD expression. This observation was replicated also in obesity (Brown et al., 2017). So far it is not known whether the link between plasticity and the susceptibility to develop addiction or obesity is innate or rather a consequence of the chronic consumption of the rewarding agent. Our data points for the first time to the possibility that the differences in synaptic metaplasticity in the VP may be an innate marker for the susceptibility to develop obesity.

It remains to unclear how a tendency to potentiate inhibitory synapses in the VP may render one more susceptible to overeat than others. A possible hypothesis is that plasticity serves as a compensation mechanism against changes in cellular activity. Thus, depressing the GABAergic input to the VP may somehow compensate for the HFHS-induced hyperpolarization in the VP. Such depression may be indeed possible in “low seekers” or some cells of OR mice (Fig. 4,5), but OP mice and “high seekers” lack this ability to weaken the inhibitory input to the VP (Fig. 4,5) and compensate for the decreased excitability. This may potentially make these latter mice more prone to develop obesity, in a yet unknown mechanism.

